# Experimental evolution of evolvability

**DOI:** 10.1101/2024.05.01.592015

**Authors:** Michael Barnett, Lena Zeller, Paul B. Rainey

## Abstract

Capacity to generate adaptive variation can evolve by natural selection. However, the idea that mutation becomes biased toward specific adaptive outcomes is controversial. Here, using experimental bacterial populations, we report the evolution of enhanced evolvability via localised hyper-mutation. Key to realisation was a lineage-level birth-death dynamic, where lineage success depended upon capacity to mutate between two phenotypic states, each optima in a cycling environment. The evolved mechanism is analogous to “contingency loci” in pathogenic bacteria, whose origin was previously unclear. Subsequent evolution showed lineages with localised hyper-mutability were more likely to acquire additional adaptive mutations. Our results provide a detailed mechanistic account of the adaptive evolution of evolvability.

## Introduction

Evolvability – the capacity to generate and maintain adaptive phenotypic variation – can be shaped by natural selection. Direct evidence comes from experimental studies of evolution in microbes where mutants with elevated global mutation rates are favoured by selection due to increased likelihood of finding beneficial variants [48, 24, 32, 51]. Evolvability in this sense is readily understood. Less clear, and often controversial, is whether (and how) selection might structure organisms so that mutation becomes biased toward adaptive outcomes [49, 41].

One reason for scepticism stems from the fact that natural selection amplifies traits beneficial in the present environment and is ‘blind’ to future contingencies [49]. Thus the suggestion that selection can structure organisms to evolve in directions *that facilitate future adaptation* is problematic. And yet, environmental change can exhibit regularities over evolutionary timescales that, in theory, might lead to biased patterns of variability [50, 56]. Indeed, *in silico* evolving populations experiencing selection toward varying and recurring goals, improve capacity to reach these goals over time [11, 27, 8, 7, 18]. This they achieve by evolving genotype-phenotype mappings in which states adaptive in the past become more readily generated by mutation. Should future selective environments resemble past ones, derived types become biased toward generating adaptive change.

A compelling case for evolvability in this sense is found in the so-called “contingency loci” of pathogenic bacteria [37]. These loci are characterised by the presence of mutation-prone sequences, such as short-sequence repeats, in genes that encode products determining interactions with the external environment. Such localised hyper-mutability causes descendant cells to achieve a diverse, yet biased set of phenotypic states, central to persistence in the face of challenges presented by host immune responses [2].

While a history of repeated encounters with a selective environment dispels any mystery of foresight, traits such as contingency loci remain difficult to explain without invoking selection at levels above the individual, which is considered weak [49, 36, 58]. However, the logic of natural selection is indifferent to level of organization or timescale, and provided there exists variation, reproduction and heredity, selection will operate [34]. In particular, selection acting on lineages provides opportunity for evolving evolvability [14, 28, 53, 19, 46, 9, 4].

## Results

We asked whether a selective regime imposed on lineages for capacity to generate adaptive phenotypic variation would lead lineages to adapt through enhanced evolvability. To this end we exploited cyclical evolutionary dynamics of the bacterium *Pseudomonas fluorescens* to provoke repeated phenotypic transitions between two niche specialist mutants [43, 42]: the mat-forming cellulose-over-producing (CEL^+^) type and the mat-colonising non-cellulose producing (CEL*^−^*) type (Figure 1). We then allowed those lineages with enhanced capacity to mutate between CEL^+^ and CEL*^−^* states to replace those lacking this ability.

**Figure 1:**
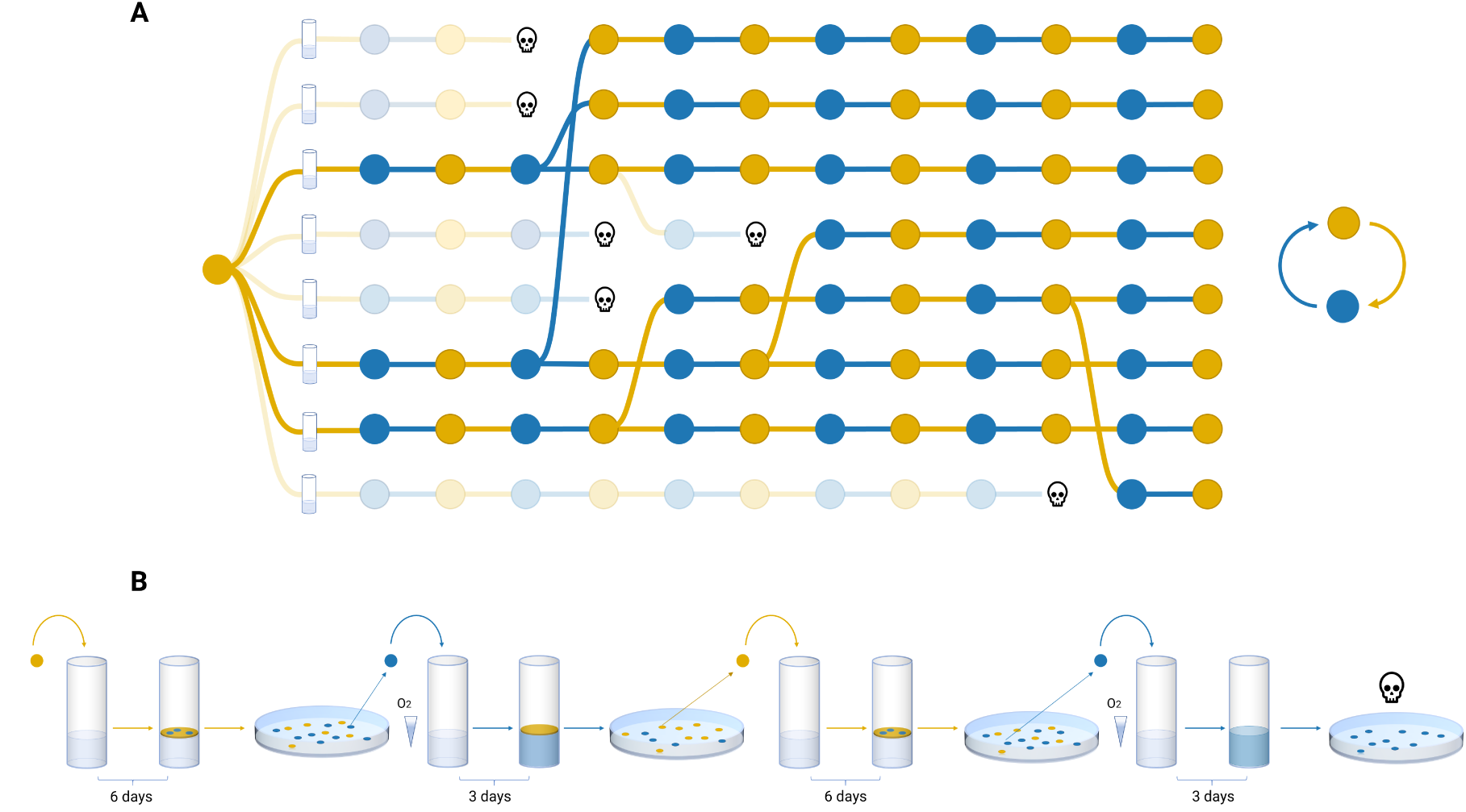
Lineage selection. **A.** Graphical representation of the experimental regime depicting eight competing lineages founded from a single common genotype. Lineages are required to transition repeatedly between CEL^+^(yellow) and CEL*^−^*(blue) states by mutation. Lineages that fail to achieve the target phenotype are marked extinct, with extinction providing opportunity for extant lineages to export their evolutionary success. **B.** Schematic of protocol that begins by founding a microcosm with a single CEL^+^ genotype. After six days in static culture, descendant cells are diluted and plated on agar with colony phenotype allowing detection of CEL*^−^*mutants. A single CEL*^−^*mutant is then used to found growth in a fresh microcosm, with a sample again examined by agar plate culture for the presence of CEL^+^ types. A single CEL^+^ type then founds the next bout of selection and the cycle repeats. Single colony bottlenecks ensure the next phenotypic state must be generated afresh via mutation. The illustrated lineage goes extinct on the final transfer because it failed to generate the next (CEL^+^) target phenotype. Key to the selection regime are frequency-dependent interactions between CEL^+^ and CEL*^−^*: CEL^+^ forms a mat at the surface and provides conditions that favour CEL*^−^* mutants; CEL*^−^* types deplete oxygen thus establishing conditions that favour the evolution of surface-colonising CEL^+^ types. Photos are shown in Figure 3E.

### Adaptive evolution of lineages

The selective regime shown in Figure 1 imposes constraints [5] that cause lineages to function as units of selection [34]. While the experiment began with a single genotype in each of the four meta-populations, mutation ensures that by the first phenotypic transition, lineages vary one to another. Failure of lineages to produce the next phenotype in the cyclical series results in extinction, with death providing opportunity for extant lineages to export their evolutionary success in a process akin to reproduction. Selection thus works over two timescales: the doubling time of cells, and the doubling time of lineages. Over the long-term, successful lineages stand to be those composed of cells that enhance lineage-level evolvability, with opportunity for cumulative refinement by selection.

Knowledge of the fate of lineages within meta-populations allows exact depiction of evolutionary dy-namics (Figure 2). Evident are selective sweeps in three of the four meta-populations (A-C) in which descendants of single lineages proliferate and form unbroken series of successful transitions. Meta-population D suffered a mass extinction at transition 48. If lineages had not been treated as units of selection, that is, if in the face of extinction, reproduction of lineages had been disallowed, then all meta-populations would have faced early extinction (Supplementary Figure 1).

**Figure 2:**
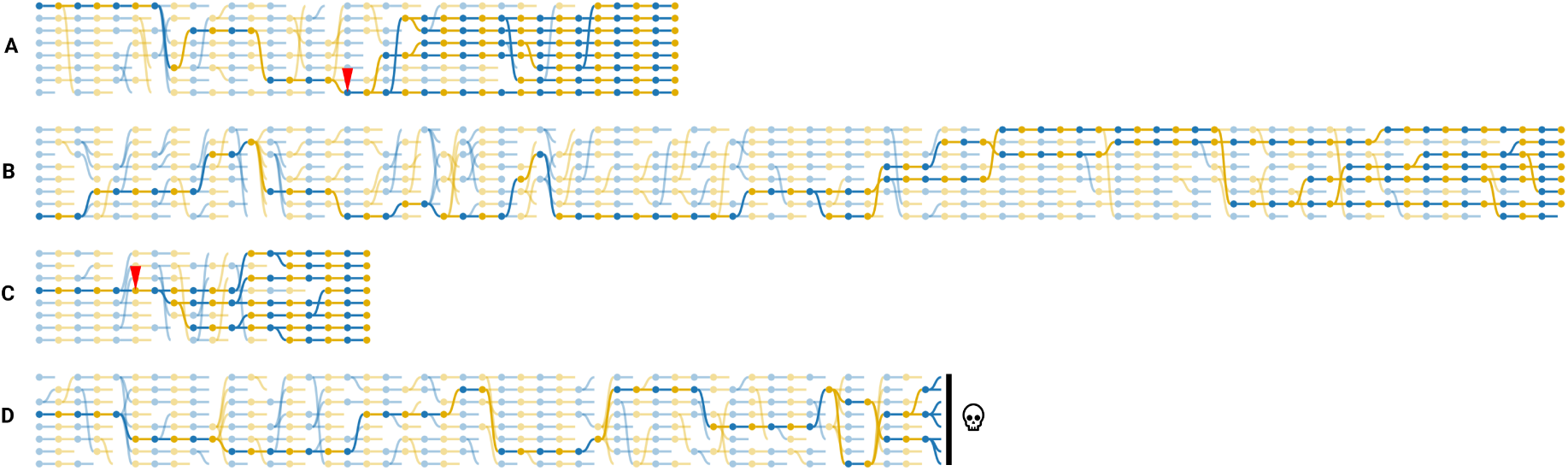
Evolutionary dynamics of meta-populations. Four CEL^+^ genotypes were used to found each of four meta-populations (A-D) each composed of eight lineages. yellow nodes indicate the CEL^+^ target phenotype and blue, CEL*^−^*. Lineage death occurred through failure to generate the target phenotype and provided extant lineages opportunity for birth. Solid lines depict the genealogy of final surviving lineages, with extinctions indicated by broken faded lines. Red arrows in A and C indicate the point of origin of mutator genotypes founding lineages that rapidly fixed (see text).

Whole-genome sequence of derived lineages from meta-populations were obtained at the end point of each series, but also periodically en route. In meta-populations A and C, derived lineages were found to carry loss-of-function (LoF) mutations in genes involved in mutational repair (*mutS* and *uvrD*, respectively), with the point of origin marked in Figure 2. Fluctuation assays confirmed global rates of mutation in meta-population A and C to be elevated 300-fold and 350-fold (Supplementary Figure 2), indicating the adaptive response of lineages was underpinned by global hyper-mutability. In contrast, no mutations affecting genes involved in DNA replication or repair were detected in meta-population B and global mutation rates remained unchanged (Supplementary Figure 2).

### Local hyper-mutability underpins lineage adaptation

Meta-population B ultimately cycled through 80 transitions (720 days within microcosms) with the experiment being terminated following stable persistence of a single lineage. To understand the genetic basis of success, whole-genome re-sequencing was used to identify and order mutations underpinning each transition between CEL^+^ and CEL*^−^*states. This revealed 548 mutational events that targeted 84 genes, 34 of which were hit multiple times. Common targets included various regulators of cyclic-di-GMP levels (394 of 548) within the cell – the intracellular signalling molecule that activates the synthesis of cellulose generating CEL^+^ – and mutations to the cellulose biosynthesis machinery itself (67 of 548). The diversity of mutational paths generated variation in evolutionary potential between lineages (Supplementary Figures S3 & S4).

Despite the availability of possible mutational paths, a single regulatory gene, *pflu0185*, became the main mutational target, eventually accounting for 59% (306 of 520) of all phenotypic transitions. Moreover, the majority of these (186 of 306) were caused by duplication or loss of a heptanucleotide (hn)-sequence ‘GGTGCCC’ located near the 5’ end of the gene (nucleotides 19-25). Mapping the occurrence of phenotypic transitions effected by changes in copy number of the hn-sequence on dy-namics of lineages within meta population B (Figure 3C) showed these mutations were associated with a decrease in the number of extinctions from 26% per transition to 10% (Figure 3A), indicating this locus underpinned lineage adaptation.

**Figure 3:**
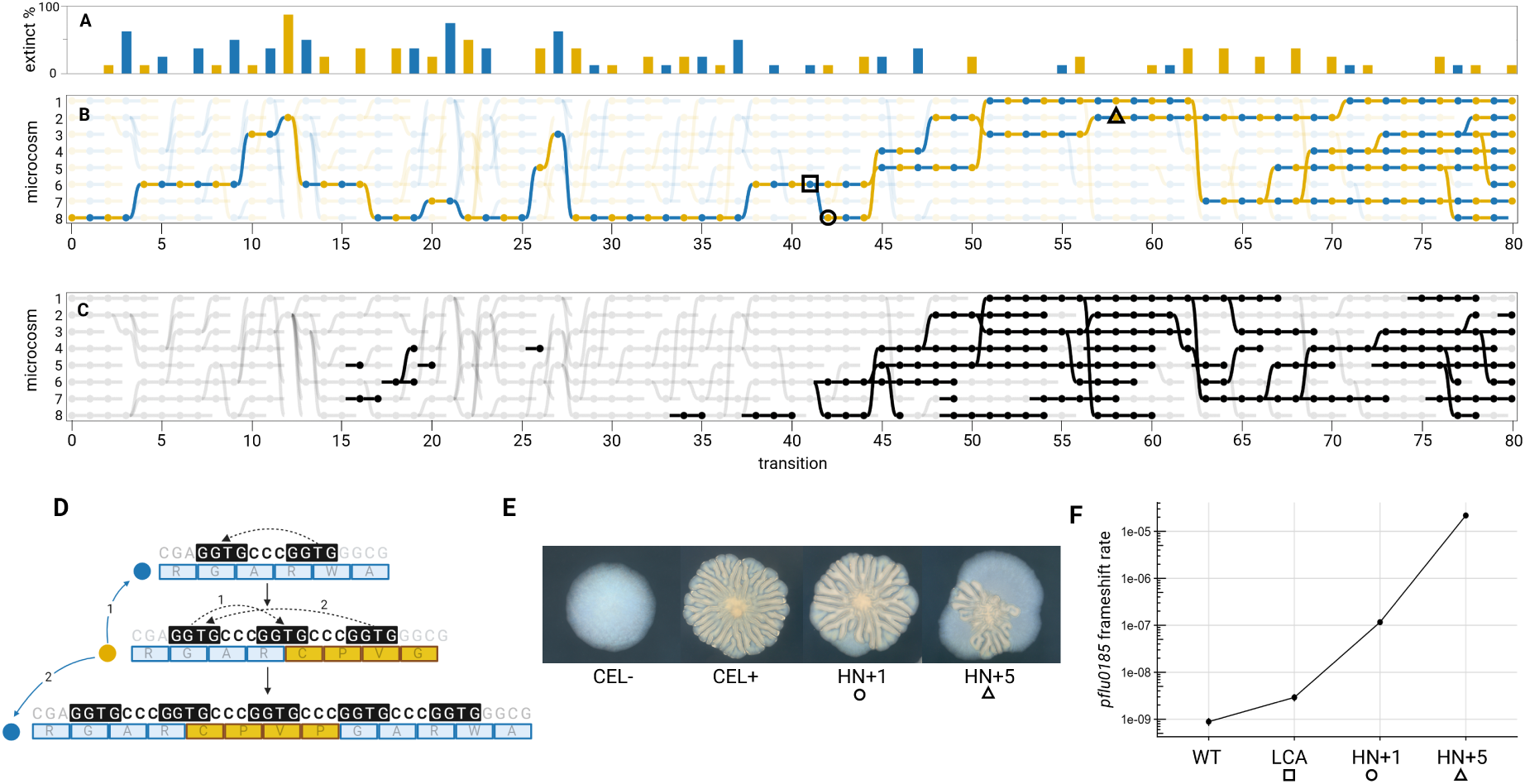
Local hyper-mutability accounts for adaptive advantage of derived lineages in meta-population B. **A**. Extinction events at CEL^+^ (yellow) and CEL*^−^* (blue) transitions. **B**. Lineage dynamics with focal points indicated by symbols around nodes (see text). X-axis labels are transitions (T) and Y-axes denotes microcosm (M). The node identified by the square box is thus T41 M6. **C**. Phenotypic transitions effected by changes in the number of copies of a hn-sequence in *pflu0185*. **D**. Proposed mechanism of slipped-strand mispairing. Top shows the DNA sequence of *pflu0185* surrounding, and including, the focal hn-sequence (GGTGCCC) plus relevant downstream nucleotides (GGTG). The dotted arrow indicates mispairing between GGTG repeats which leads to an additional copy of the hn-sequence (middle, vertical arrow). With two copies of the hn-sequence the reading frame is destroyed. The reading frame can be corrected by either mispairing in the forward direction (dotted arrow 1), which restores the original state (blue arrow 1); or, mispairing in the reverse direction (dotted arrow 2) resulting in four copies of the hn-sequence (bottom, vertical arrow). **E**. Image of CEL*^−^* and CEL^+^ colonies following prolonged (72 h) growth on agar plates with derived lineages having either a single additional copy of the hn-sequence, or five additional copies, the latter showing overt sectoring indicative of an elevated mutation rate. **F**. Relationship between genotype and rate of frame-shift mutations in *pflu0185* determined by fluctuation assay. Symbols denote genotypes from (B). Error bars (largely contained within the points) are 95% CI.

Mechanistically, rapid changes in the number of copies of short sequence repeats can be attributed to slipped-strand mispairing — a genomic instability that increases the likelihood of deleting or duplicat-ing repeated units [33]. Relevant here is that downstream of the duplicated hn-sequence (GGTGCCC) in ancestral *pflu0185* is a further copy of GGTG and thus the focal sequence is GGTGcccGGTG (Figure 3D). Mis-pairing between the two copies of GGTG stand to catalyse an initial hn duplication event. The resulting addition of the hn-sequence disrupts the *pflu0185* reading-frame near its N-terminus, thereby eliminating protein function.

Pflu0185 is a regulatory protein composed of two enzymatic domains: a diguanylate cyclase (DGC) — responsible for the synthesis of cyclic-di-GMP — and a phosphodiesterase (PDE) — responsible for its breakdown [45]. In *P. aeruginosa* functional analysis shows the protein to have PDE function, but no DGC activity [6]. This is consistent with the genetic analysis reported here. As a consequence of the initial hn duplication, PDE function is lost and cyclic-di-GMP levels increase within the cell. Elevated levels of cyclic-di-GMP in turn activate the cellulose biosynthesis machinery, and cells enter the CEL^+^ state.

Beyond the adaptive CEL^+^phenotype, cells inheriting the initial hn duplication in *pflu0185* also inherited an enhanced ability to generate the CEL*^−^*phenotype, being *∼*five times more likely to survive the next transition compared to inheritance of any other mutation (95% CI [1.8, 12.9], *P* =0.0006, Fisher’s exact test, two-tailed). This increased survival can be explained by the relationship between number of tandem repeats and rate of mispairing, where a rise in the number of repeats is expected to elevate local mutation rate [2]. The effect is that cells with the hn duplication are primed to re-activate *pflu0185* function through indels that restore the reading frame. Indeed, 96% of lineages inheriting the initial duplication corrected the reading frame through subsequent expansion or contraction of the hn-repeat. In 86 of 89 instances this involved restoration of the reading frame by contraction. In three instances the reading frame was restored by expansion of the tract through addition of two hn copies. However, lineages that expanded the tract were then more likely to see further expansion in copy number of the hn-sequence at the next transition (5:3 contraction:expansion, *P* =0.0049, Fisher’s exact test, two-tailed). As the number of repeats increased, the rate at which transitions occurred visibly increased (Figure 3E).

While global measures of mutation rate reported above showed no increase, by fusing *pflu0185* to an out-of-frame kanamycin resistance (*kanR*) gene, the rate of frame-shifting, specifically within *pflu0185*, could be quantified by fluctuation assay (Figure 3F). A statistically significant three-fold increase in the rate of frame-shifting was evident in the genotype that marked the coalescent point of the final lineages (T41 M6), black square in Figure 3B), despite the fact that this genotype carried no additional copy of the hn-sequence. With two copies ((T42 M8), black circle in Figure 3B), the rate of frame-shifting increased a further 150-fold, and with six (+5) copies (T58 M2, black triangle in Figure 3B) (the maximum observed in the selection experiment) the rate of frame-shifting was more than four orders of magnitude greater than the ancestral genotype (Figure 3F).

### Concomitant increases in transcription and mutation rate at *pflu0185* potentiated repeat-associated hyper-mutation

Having established localised hyper-mutability as underpinning lineage adaptation in meta-population B, we next sought understanding of evolutionary origins. In particular, questions remained as to why initial duplication of the hn-sequence had become increasingly probable, despite any LoF within *pflu0185* being capable of achieving the same effect. Indeed, alternative LoF mutations in *pflu0185* were still occasionally observed late in evolution and, where irreparable, led to lineage extinction (Supplementary Figure 5).

Focus therefore turned to the series of mutations targeting *pflu0185* that occurred prior to the expansion of lineages at T41 M6. By this point *pflu0185* had acquired 24 mutations (including three direct reversals), either within, or upstream of the gene, with each mutation being responsible for the transition between CEL^+^ and CEL*^−^* states. Among this convoluted history, no single event stood out in terms of relevance to hyper-mutability caused by changes in copy number of the hn-sequence. However, given the presence of multiple mutations in the upstream region — some of which led to formation of recognisable promoter motifs (Figure 4B) — and recognising the mutagenic nature of transcription [13], attention turned to the dual effects of upstream mutations on levels of transcription and rate of frame-shifting in *pflu0185*.

**Figure 4:**
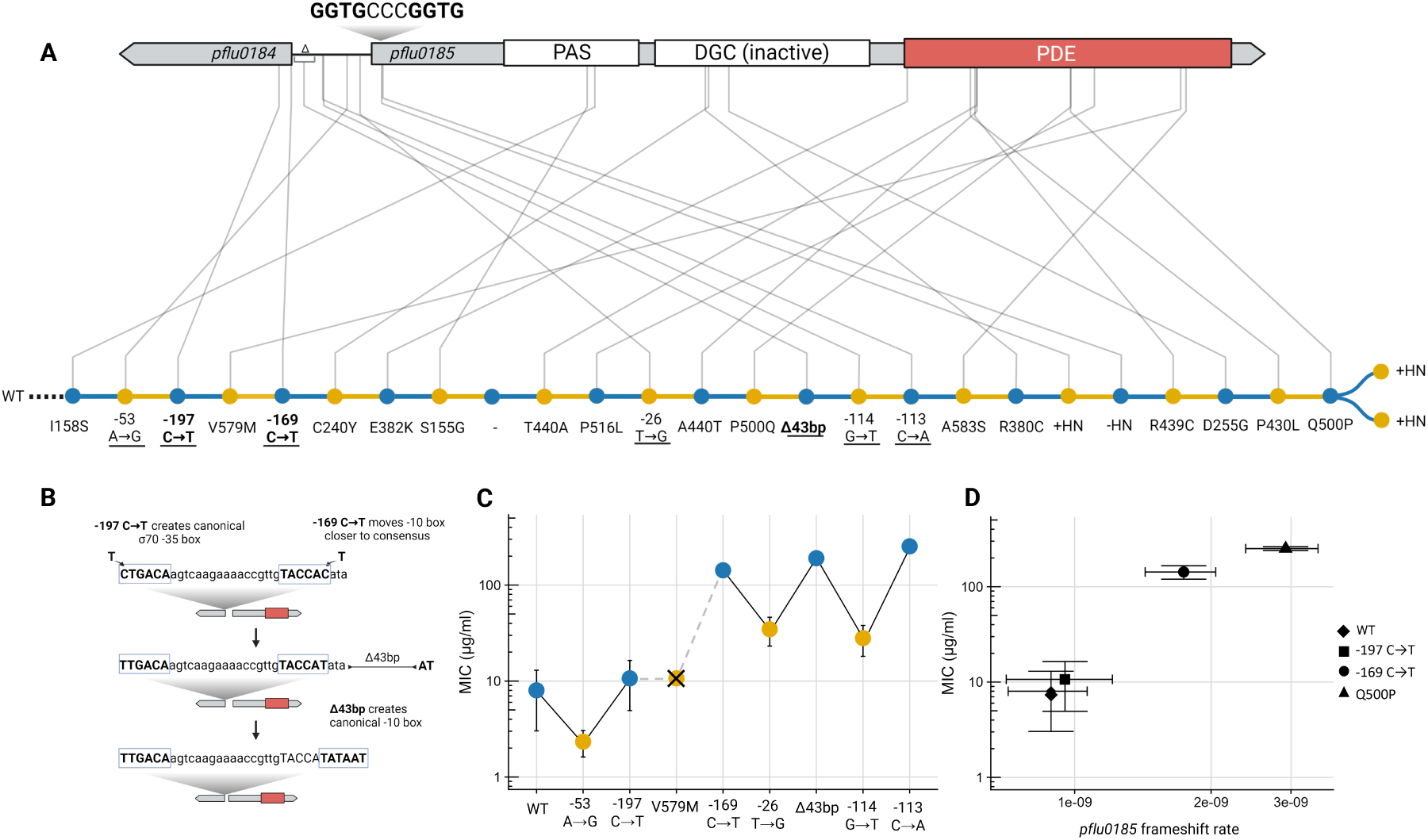
Mutational history of potentiating mutations in *pflu0185*. **A.** Genetic structure of *pflu0185* and sequence of successive mutations modulating changes in phenotypic state. The top portion of A shows the *pflu0185* open reading frame with location of the hn-sequence, PAS signalling domain, inactive DGC domain and, in red, the PDE domain. Also shown is the upstream gene and intergenic space where multiple mutations arose that affected transcription. **B**. Effects of mutations on transcription of *pflu0185* and CEL^+^/ CEL*^−^* status determined through assay of MIC in the presence of kanamycin (putative promoter mutations are underlined in A). Mutation V579M is included to demonstrate its role in reducing catalytic activity of the PDE, necessitating further increases in *pflu0185* transcription to trigger the CEL*^−^* state. **C**. Mutational changes in upstream regions (in bold type) placed in context of surrounding DNA sequence that contribute to formation of a typical *Pseudomonas* promoter.[16, 52] **D**. Relationship between transcription (as determined by MIC assay) and rate of frame-shift mutations in *pflu0185*. Note that the MIC of Q500P is equivalent to the last upstream mutation (*−*133 C→A). Error bars are 95% CIs.

To assess effects of mutations on *pflu0185* transcription, we took advantage of the *pflu0185-kanR* gene fusion, using level of kanamycin resistance (expressed as minimum inhibitory concentration (MIC)) as a proxy. This revealed a ratcheting effect of mutations, with those causing either a reduction in transcription of *pflu0185*, or reduction in catalytic activity of the PDE (leading to the CEL^+^ state), being compensated for by mutations in the upstream region that elevated transcription (leading to the CEL*^−^* state) (Figure 4C). By the end of this series of mutations, transcription of *pflu0185* was elevated 30-fold above that of ancestral *P. fluorescens*.

In terms of the possibility that these mutations had potentiated proclivity of the hn-sequence to expand in copy number, the relationship between MIC and rate of frameshift mutations in *pflu0185* was determined by fluctuation assay. This revealed a clear positive correlation (Figure 4D). Notably, a promoter mutation activating *pflu0185* also preceded the first instances of mutation via the hn-sequence, which occurred independently in a sub-lineage that later went extinct (T16 M5 and M7 (Figure 3C)).

### Stringent selection drives expansion in HN copy number and stabilizes local hyper-mutability

Despite evident potential for self-accelerating increases in mutation rate by successive duplication of repeats, all lineages that expanded the hn-sequence periodically reverted to a single copy. This meant lineages often relied on the relatively weak rate of initial hn duplication for survival. Consequently, extinction events persisted, due primarily to transitions mediated by irreparable destruction of the cellulose biosynthesis machinery (Supplementary Figure 4).

We reasoned that this tendency was in part a consequence of high mutation supply within microcosms (a product of population size and propagation time), which meant extreme mutation rates (Figure 3F) were not necessary to reliably generate cells with the target phenotype. Indeed, contingency loci found in nature often consist of many tandem repeats [2], suggesting more stringent lineage-level selection pressures. Accordingly, we took a single CEL^+^ genotype carrying the initial hn-duplication (+1 GGTGCCC unit) (T42 M8 black circle in Figure 3B) and used this genotype to found 32 lineages that competed under a restrictive regime where population size was reduced by a factor of thirty and where survival required generation of the target states every 24 h (Figure 5).

**Figure 5:**
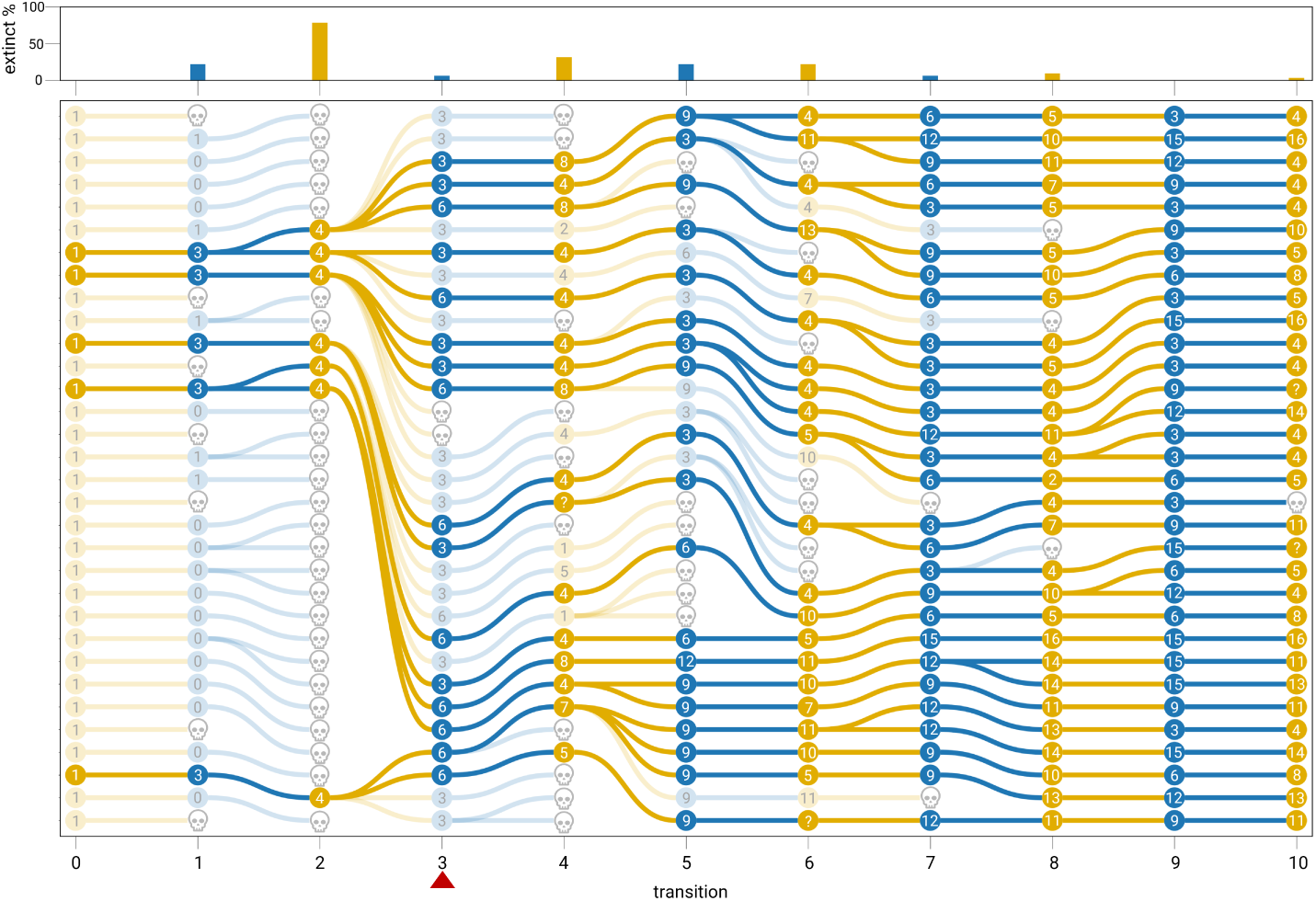
Stringent selection drives expansion in HN copy number. A total of 32 lineages were founded by genotype T42 M8 (black circle in Figure 3B) containing a single additional copy of the hn-sequence. Lineages were subject to the selection regime as shown in Figure 1, but population size was reduced 30-fold and lineages were required to achieve the target phenotype within 24 hours or face extinction. The number of additional hn units over the ancestral SBW25 *pflu0185* sequence is indicated within nodes. Colour code for phenotypic states are as described previously. The red triangle indicates a point after which the stringency of selection was further increased by requiring successful lineages to have at least five colonies of the target phenotype present. The upper panel shows extinctions at each transition.

The selective response, lineage-level dynamics, and changes in numbers of repeats over ten transitions are revealing. Most lineages managed — within 24 h — to achieve the CEL*^−^* state required for the first transition, but the majority did so by losing one copy of the hn-sequence. Without exception, those lineages that switched to CEL*^−^* by loss, were unable to transition to the CEL^+^ state during the next 24 h period and thus went extinct. In sharp contrast, lineages that achieved the CEL*^−^* state (at transition 1) by adding three additional copies, became progenitors of lineages that persisted throughout the course of the experiment. At the end point, lineages harboured on average 8 additional copies of the hn-sequence, with some acquiring as many as 16. A clear association is evident between the number of copies of the hn-sequence and likelihood of survival.

### Localised hyper-mutability of *pflu0185* -facilitated additional adaptive evolution

A final observation concerns transitions in the initial experiment associated with multiple mutations. Specifically, we noted that during the 6-day CEL^+^to CEL*^−^*period, transitions effected by changes in hn-repeat number were more than three times as likely to be associated with additional (secondary) mutations. Of 91 such transitions, 19 harboured secondary mutations (21%). In contrast, of 167 transitions effected by mutations not involving the hn-sequence, only 10 carried secondary mutations (6%) (*P* =0.0007, Fisher’s exact test, two-tailed).

While secondary mutations that hitchhike with transition-causing mutations are to be expected (Supplementary Table 2), mutations in loci encoding genes connected to chemotaxis and motility (Supplementary Table 3), have conceivable ecological relevance. Moreover, two loci, the aerotaxis receptor *aer* (*pflu4551*) and a methyl-accepting chemotaxis protein (*pflu1687*), were observed to occur repeatedly (six and four times, respectively) across multiple lineages during transitions mediated by the hn-sequence (Figure 6A). Such striking parallelism suggested these mutations were themselves adap-tive. Figure 6B elaborates the benefit presumed to accrue to lineages that switch phenotype early, and reliably.

**Figure 6:**
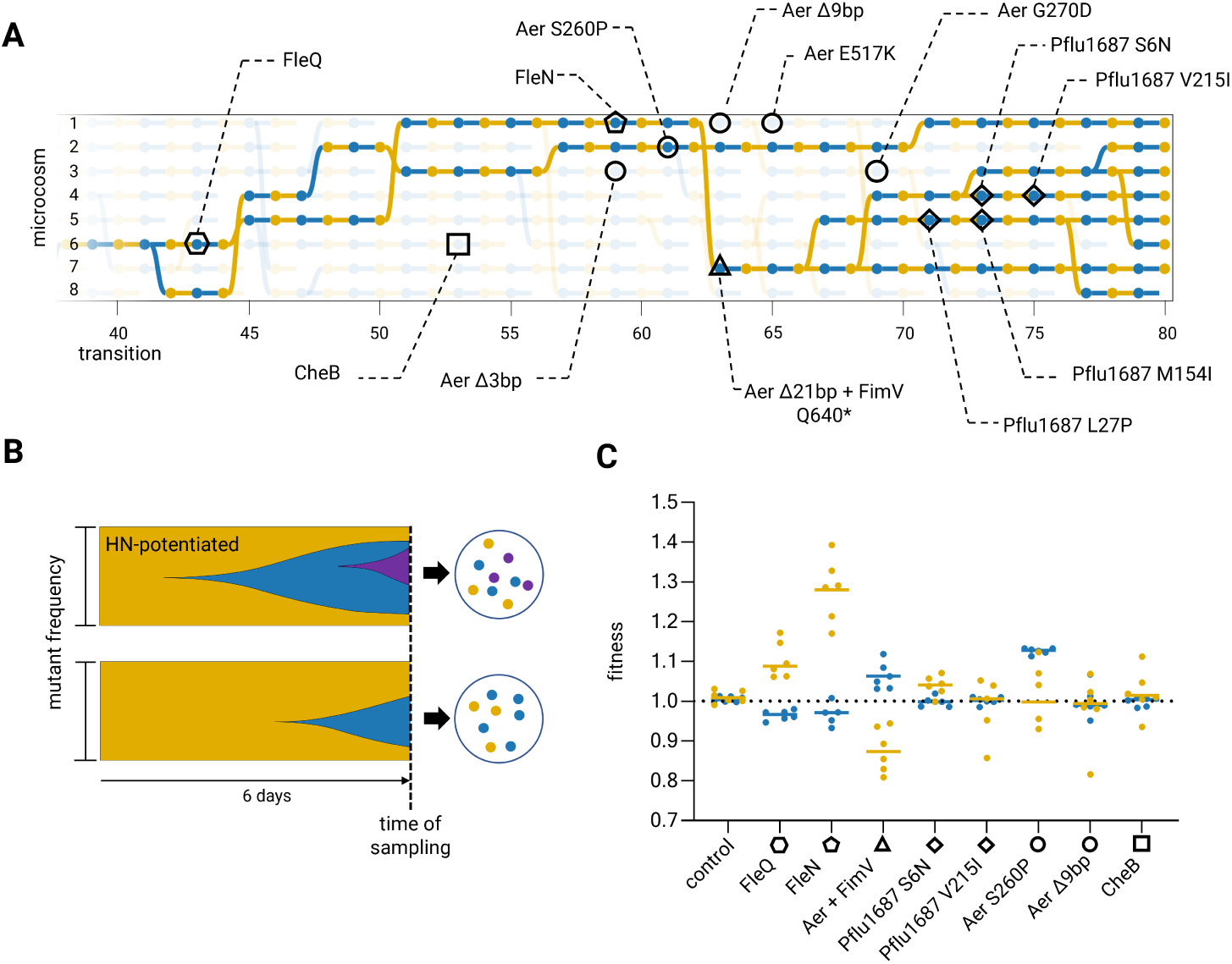
Secondary mutations and fitness effects. **A**. Genealogy of successful lineages from meta-population B between transition 40 and 80 on which secondary mutations of predicted ecological significance are indicated. **B**. Muller plots showing predicted consequences of presence (top) and absence (bottom) of local hyper-mutation: early and reliable switching provides enhanced opportunity for secondary mutations (purple) to reach high frequency (top). **C**. Fitness consequences of mutations determined by competitive assay against immediate ancestral types in CEL^+^(yellow) and CEL*^−^*(blue) backgrounds. Lines indicate mean value.

To test whether these secondary mutations were adaptive, the fitness effects of a sample of mutations were determined relative to immediate ancestral types, both in CEL^+^ and CEL*^−^* backgrounds (Figure 6C). Following the lineage that descends from T42 M6, the first mutation examined (in T43 M6) was in FleQ (R371C), a master regulator of motility and polysaccharide production (including cellulose) [25, 21]. Relative to the immediate ancestral types the *fleQ* mutation increased fitness *∼*10% in CEL^+^, but marginally decreased fitness in CEL*^−^*. The next mutation, in FleN (L42Q), which also regulates flagella through interactions with FleQ [12], increased fitness of CEL^+^ by a further *∼*30%, with minimal effects in CEL*^−^*. Relative to the fitness of T42 M6, the two mutations combined contributed a *∼*40% fitness gain in CEL^+^.

The genotype at T62 M7 carries two additional mutations (one of only three instances in the experiment where three mutations co-occurred at the same transition), one a 21 base pair deletion in Aer and the other a stop codon within an additional motility regulator FimV (Q640*). These two mutations reduced fitness of CEL^+^ by *∼*10%, while increasing fitness of CEL*^−^*by *∼*5%. With the exception of Pflu1687 (S6N) in the CEL^+^ state, no further fitness effects were detected. Secondary mutations in other lineages were also examined, including changes in the chemotaxis protein CheB (T92P) and further mutations in Aer (S260P and Δ9bp), with the largest contribution coming from Aer S260P in the CEL*^−^* state). Together the range of effects shown in Figure 7C are consistent with the hypothesis that at least some secondary mutations co-occurring with changes in hn-repeat number are adaptive.

## Discussion

Evolution by natural selection is a blind process, but living systems can appear to possess evolutionary foresight [14, 26]. Mechanistically, this is conceivable. Certain configurations of gene regulatory networks [55, 11, 56], developmental systems [28], chromosomal architectures [8, 40], and mutational processes [37, 22] have apparent adaptive utility in future environments. Ability to realise such advantages requires not only memory of past history, but often an ability to regenerate previously achieved phenotypic states. Here we have detailed how selection on lineages can incorporate prior evolutionary history into the genetic architecture of a single cell such that mutation appears to anticipate future environmental change.

Central to our findings was a selective process where lineages better able to generate, by mutation, adaptive phenotypic variants, replaced those that were less proficient (Figure 1). In one metapopulation, a single lineage emerged with capacity to transition rapidly between phenotypic states via expansion and contraction of a short nucleotide repeat in a manner precisely analogous to that of contingency loci in pathogenic bacteria [37]. Moreover, when multiple lineages of a single genotype carrying the mutational history of 42 previous transitions, but carrying just a single additional copy of the hn-sequence, were propagated under a regime that required phenotypic switching on a daily basis, lineages with up to 16 copies of the repeat rapidly evolved (Figure 4). The derived locus is thus itself highly evolvable with respect to alterations in the frequency of environmental change.

Selection works most forcefully on individuals and is typically weak at higher levels of organisation [36, 58, 49]. In our experiments, the death and birth of lineages established a feedback between lineage-level properties and the evolutionary dynamics of cells. While selection within microcosms favours fast-growing cells, successful lineages were those with capacity to generate phenotypic variants suited to the future environment. Selection over the doubling time of lineages served to align cell and lineage fitness, precisely as occurs during, for example, the evolutionary transition from cells to multicellular life [5].

Adding potency to lineage selection was the fact that, throughout our experiment, variation in evolutionary potential between lineages resulted from how adaptive CEL^+^ and CEL*^−^* phenotypes were instantiated by mutation. That a particular history of mutations — each selected due to its immediate benefit to the cell — could culminate in the variation necessary for construction and refinement of a lineage-level adaptation was essential for aligning cell and lineage fitness. This departs from the common assumption — and basis for doubting the efficacy of selection on evolvability — that mutations altering evolutionary potential deliver no immediate benefit to the organism and are therefore susceptible to loss via drift [36, 49].

The interplay between mutations conferring immediate benefit to cells and evolvability over the longterm [54] is best illustrated by the convoluted mutational history of *pflu0185* that preceded its hyper-mutator status. Critical to obtaining this status were a series of mutations leading to elevated *pflu0185* transcription. Each mutation increasing *pflu0185* transcription was selected within microcosms on the basis of realisation of the adaptive CEL*^−^* state. In particular, promoter mutations compensating previous intragenic LoF mutations that impaired catalytic activity of Pflu0185 were crucial in ratcheting increased transcription. Elevated transcription then indirectly potentiated duplication of the hn-sequence, consistent with previous observations of transcription-associated mutagenesis [57, 60].

Evidence of additional (secondary) adaptive mutations associated with local hyper-mutation of *pflu0185* led us to reason that increased propensity to switch enhances the possibility that secondary adaptive mutations, which would otherwise be stalled by the waiting time required to achieve the primary transition-mediating mutation, can increase in frequency (Figure 6B). Data presented in Figure 6 supports this hypothesis, showing how the speed of evolution can increase without the threat of Muller’s ratchet that arises from global hyper-mutation [47, 10]: ability to generate adaptive phenotypic variation provides a ‘head-start’ to adapt to other aspects of the local environment with feedback affecting further lineage success.

Finally, we turn to broader implications of the study and particularly the extent to which findings here might have general relevance. The selective regime employed was contrived, with selection on lineages being strictly enforced. Such stringent conditions are likely limited in nature. However, microbial pathogens faced with the challenge of persistence in face of the host immune response, will experience strong lineage-level selection, with repeated transitions through selective bottlenecks [38]. As we have shown here, precisely these conditions can promote the evolution of evolvability. That the genomes of pathogenic bacteria are replete with contingency loci bears testimony to this fact. Notably, while each such locus is likely constructed through repeated encounters with specific selective environments, possession of multiple loci enables combinations of phenotypic states that stand to enhance evolvability in novel scenarios (cf. [39]).

But we further suggest that selection on lineages is more important than generally appreciated [23]. Lineages need only differ in evolutionary potential, for example in how adaptive phenotypes are instantiated by mutation. Provided they co-exist for sufficient time that phenotypic consequences manifest, then lineages differing in capacity to generate adaptive variation stand to compete [59, 20, 31]. This will be particularly relevant in situations where transmission / migration involves passage through bottlenecks that require variation to be generated afresh [29, 35, 3]. Moreover, as demonstrated here, if challenges to lineage survival repeat across time, then adaptations that enhance evolvability — shaped by lineage selection for the benefit of lineages — are to be expected.

## Materials and Methods

### Experimental evolution

Both CEL^+^ and CEL*^−^* were propagated 25 ml glass microcosms containing 6 ml of King’s Medium B (KB) at 28°C, without shaking. The four meta-populations were founded by single genotypes derived from a previous, but conceptually similar, experiment [24]. Founding genotypes carried on average nine unique mutations, primarily in characterized c-di-GMP regulators (Supplementary table 4).

For the CEL^+^ to CEL*^−^* phase, microcosms were incubated for six days, vortexed, and 50 *µ*l of a 3.35 × 10^5^-fold dilution spread on a KB agar plate. After 48 hours growth each plate was then examined for the presence of CEL*^−^* colonies, with a single colony used to inoculate a fresh microcosm and initiate a three day growth period. 100 *µ*l of an 8.0 × 10^6^-fold dilution of each microcosm was plated and screened for CEL^+^ types. The process was iterated. A random number generator was used to determine the extant lineage that would replace extinct lineages. When an entire meta-population failed to generate the target state twice over, propagation was ceased (as in meta-population D).

The births and deaths of lineages were recorded, and the resulting lineage dynamics visualized using Colgen [17]. Overnight cultures of each selected CEL*^−^*and CEL^+^ colony were stored at -80°C in 33% glycerol saline. The ‘stringent selection’ experiment followed the same procedure as above but with cells evolved in 96-well plates containing 200*µ*l of KB, and a reduced period of 24 hours given before dilution and screening.

### Sequencing and detection of mutations

Visualization and knowledge of genealogy was used to inform both whole-genome re-sequencing and targeted Sanger sequencing to detect and order mutational events. For meta-population B, 482 clones representing successive time-points of the experiment were sequenced, allowing precise ordering of mutations underpinning the majority of the 520 transitions. Remaining transitions and associated mutations were then inferred by parsimony or confirmed by Sanger sequencing. In one instance, to resolve an ambiguity, a mutation (-197 C→T *pflu1085*) was re-constructed to confirm its phenotypic effect.

Genomic DNA was extracted from overnight cultures using the DNeasy DNA Blood & Tissue Kit (Qiagen) according to the manufacturer’s spin column protocol. Library preparation and whole-genome sequencing to a depth of *∼*50x coverage was conducted at either the Max Planck Institute for Evolutionary Biology (Plön, Germany) or Eurofins Genomics (Constance, Germany) using Illumina MiSeq, NextSeq, or NovaSeq. The resulting reads were assembled against the reference *P. fluorescens*SBW25 genome (GenBank accession:AM181176.4) using the clonal option of Breseq (version 0.35.5) with default parameters [15]. Mutations were identified with breseq’s HTML output and in the case of predicted novel junctions manually inspected with Geneious Prime. Marginal predictions were confirmed by Sanger sequencing. PCR products for Sanger sequencing were purified with EXOSAP and sequencing conducted by the Max Planck Insitute for Evolutionary biology, prepared using the BigDye Terminator v3.1 cycle sequencing kit followed by the BigDye XTerminator purification kit (ThermoFisher) according to manufacturer’s instructions.

### Strain construction

Isothermal Gibson Assembly reactions with the NEBuilder HiFi DNA Assembly master mix from New England BioLabs, were used for all molecular cloning procedures. Constructs to be integrated into the SBW25 genome contained flanking regions of 400-900 bp homologous to the site of integration and were assembled into the transfer plasmid pUISacB (accession pending). Primer overhangs were generated using the online NEBuilder tool ( nebuilder.neb.com) and are available on request. All PCRs were performed using master mixes of high-fidelity polymerases (Phusion or Q5) and purified with the QIAquick PCR purification kit (Qiagen). pUISacB was extracted from overnight cultures using the QIAprep Spin Miniprep Kit (Qiagen), linearized with the restriction enzyme SpeI, amplified by PCR and digested with DpnI to remove remaining plasmid template. Assembled constructs were transformed into chemically competent *E. coli* Top 10 cells and integrated into SBW25 via two-step allelic exchange as previously described [1], but with final mutant selection using sucrose counter-selection on tryptone yeast extract and sucrose (TYS10) plates (10 g/L tryptone, 5 g/L yeast extract, 10% (w/v) filtered sucrose, 15 g/L agar).

For the *pflu0185* ‘frame-shift detector’ used to measure local frame-shift rate, a fusion was constructed with *kanR* via the linker sequence ’GCTAGCAGCGCAAGCGCAAGCGCA’ (ASSASASA), such that the latter gene was 1bp out of frame with respect to *pflu0185* (an in-frame version of the same *kanR* fusion was also constructed). Three synonymous mutations engineered into the 5’ end of *kanR* were required to remove aberrant start codons that existed in the out-of-frame sequence (which otherwise re-started the correct reading-frame). Strains expressing fluorescent proteins for use in fitness assays were constructed via standard conjugation of recipient SBW25 cells and *E. coli* donor containing pMRE-Tn*7* -145 (mScarlet-I) or pMRE-Tn*7* -152 (sGFP2) with media containing 0.1% w/v arabinose to induce Tn*7* transposase genes [44].

### Transposon mutagenesis

Transposon mutagensis using Tn*5* was conducted according to the previously described method [21]. Briefly, this involved a tri-parental conjugation between the recipient CEL^+^ and an *E. coli* donor containing the plasmid pCM639 with the IS-Ω-kan/hah transposon along with *E. coli* containing the helper plasmid pRK2013. Suppressor mutants were identified by screening for CEL*^−^* colonies. The site of insertion was determined by amplifying the transposon-chromosome junction through arbitrarily-primed PCR with Sanger sequencing of the resulting product.

### Minimum inhibitory concentrations

Minimum inhibitory concentration (MIC), used as a proxy for transcription, was determined by plating *∼*10^8^ cells of the relevant genotypes engineered with an in-frame *pflu0185* :*kanR* fusion onto KB agar and applying a kanamycin MIC test strip (Liofilchem). After 24 hours growth the MIC was determined by identifying where the margin of growth intercepted the strip in three independent replicates. In cases where genotypes exhibited high levels of resistance extending beyond the maximum measurable range of the test strips (256 *µ*g ml^-1^), the assay was instead conducted on agar containing either 100 or 200 *µ*g ml^-1^ of kanamycin and the concentrations summed.

### Fluctuation assays

Colony counts on selective and non-selective plates were used to estimate mutation rates using the rSalvador R package (v1.9), which generates a maximum likelihood estimate of mutation rate under the Luria-Delbruck model while accounting for variation in plating efficiency[61]. The likelihood ratio method was used to calculate confidence intervals and the statistical significance of differences between mutation rate estimates. Global mutation rates were assessed using rifampicin (100 *µ*g ml^-1^) and kanamycin (at concentrations of 10-50 *µ*g ml^-1^ depending on *pflu0185* expression) used to determine rates of frame-shifting within *pflu0185*. Twelve independent cultures were used to determine spontaneous mutants; 6 were used to determine population size. To minimize confounding fitness differences between CEL*^−^* and CEL^+^ states, all strains used for *pflu0185* frame-shifting fluctuation assays were modified with a cellulose synthase inactivating mutation (WssEΔ993L-1260E) so that cells maintained the CEL*^−^* state.

### Fitness assays

Fitness was measured by 1:1 competition in either static (for CEL^+^) or shaken (for CEL*^−^*) microcosms. Shaking microcosms were used as a proxy environment for CEL*^−^* due to difficulty of reconstituting the selective conditions of growth within a CEL^+^ mat. Competitors were fluorescently marked with either mScarlet-I or sGFP2 and were otherwise isogenic but for the secondary mutation of interest. The control consisted of a single representative CEL*^−^* or CEL^+^ strain with both fluorescent markers competed against itself. Starting populations were derived from colonies inoculated into KB microcosms and grown 24 hours at 28°C in a shaking incubator. Microcosms were then vortexed and in the case of CEL^+^ strains, sonicated (Bandelin Sonopuls mini 20; 2mm probe, 80% amplitude, 20 second duration) to further break up cell aggregates. Competitors were then mixed 1:1 and diluted in filtered PBS (0.22*µ*M filter) with the initial ratio of fluorescent events obtained using flow cytometry (MACSQuant VYB). 6 *µ*l of the 1:1 mix was also used to initiate competitions, which ran for either 24 (shaken) or 48 (static) hours. Following competitive growth, microcosms were vortexed (and static CEL^+^ microcosms again sonicated), diluted in filtered PBS, and the final ratios determined by flow cytometry. Fitness was calculated as the ratio of Malthusian parameters [30].

## Supporting information

Supplementary video 1 - mutational trajectories

## Acknowledgements

The authors acknowledge generous core support from the Max Planck Society and thank members of the Department of Microbial Population Biology for help, ideas and discussion, and Alina Waldmann for genomic DNA purification. The authors are especially grateful to David Rogers for insight into the mechanism by which the hn repeat achieves the first duplication and suggestion of a strategy for measuring frame-shift mutations. We are also indebted to Richard Moxon for valuable comment and discussion.

## Author contributions

MB and PBR: experimental concept and design. MB: experimental work and analysis with technical assistance by LZ. MB and PBR wrote the paper.

## Data availability

All data are available in the manuscript, Supplementary Information or retrievable from Edmond: https://doi.org/10.17617/3.5PEE52. All genetic data and the full history of meta-population B in the form of an interactive tree can be explored at https://micropop.evolbio.mpg.de/data/evolve

