## Supplementary Information for "Experimental evolution of evolvability"

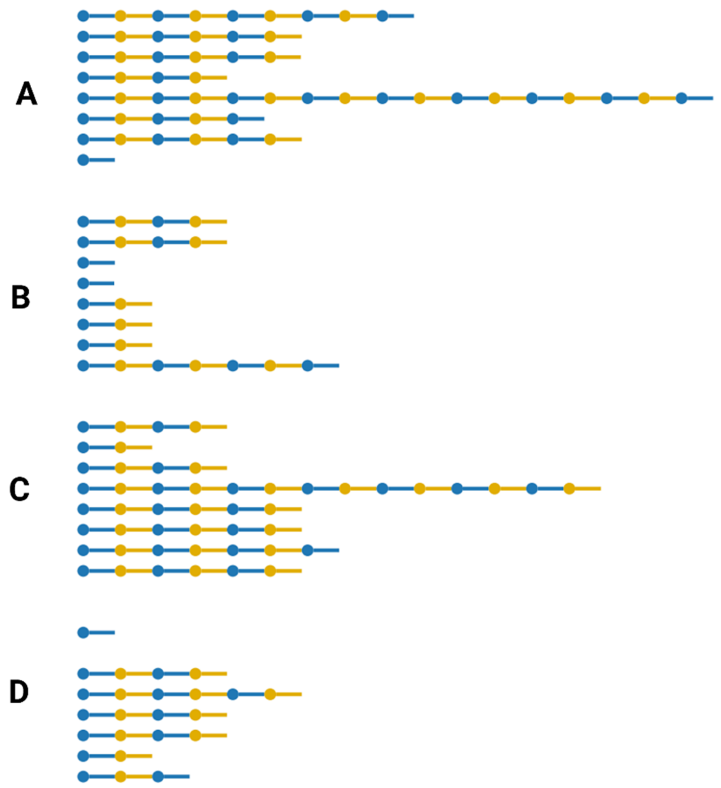


**Figure S1. Lineage dynamics in the absence of reproduction.** The genealogies are from meta-populations A-D shown in Figure 2, but here the effects of disallowing lineage reproduction are shown. The difference between lineage data here, versus that displayed in Figure 2, is attributable to the effect of selection acting on viability and fecundity (as in Fig. 2) versus viability alone. In the absence of reproduction there is limited chance that selection can cumulatively shape the evolution of evolvability.

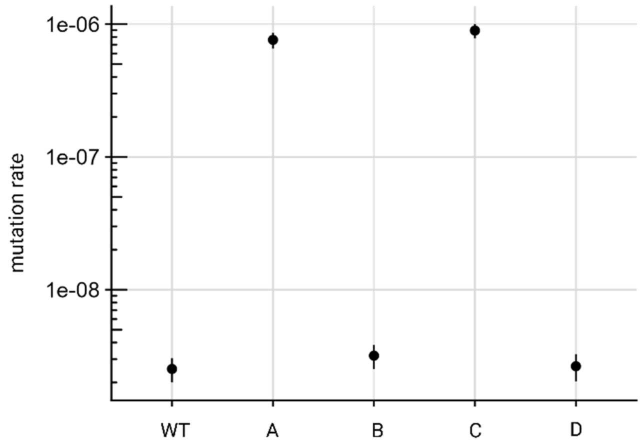


**Figure S2. Global mutation rate estimates for ancestral (wildtype) *P. fluorescens* SBW25 and experimental genotypes derived from surviving lineages in meta-populations A, B, C and D.** Mutation rate is the number of mutations per cell replication conferring rifampicin resistance. Errors bars represent 95% CI. The difference between wildtype and the representative genotype from meta-population B was not significant (*P*=0.12).

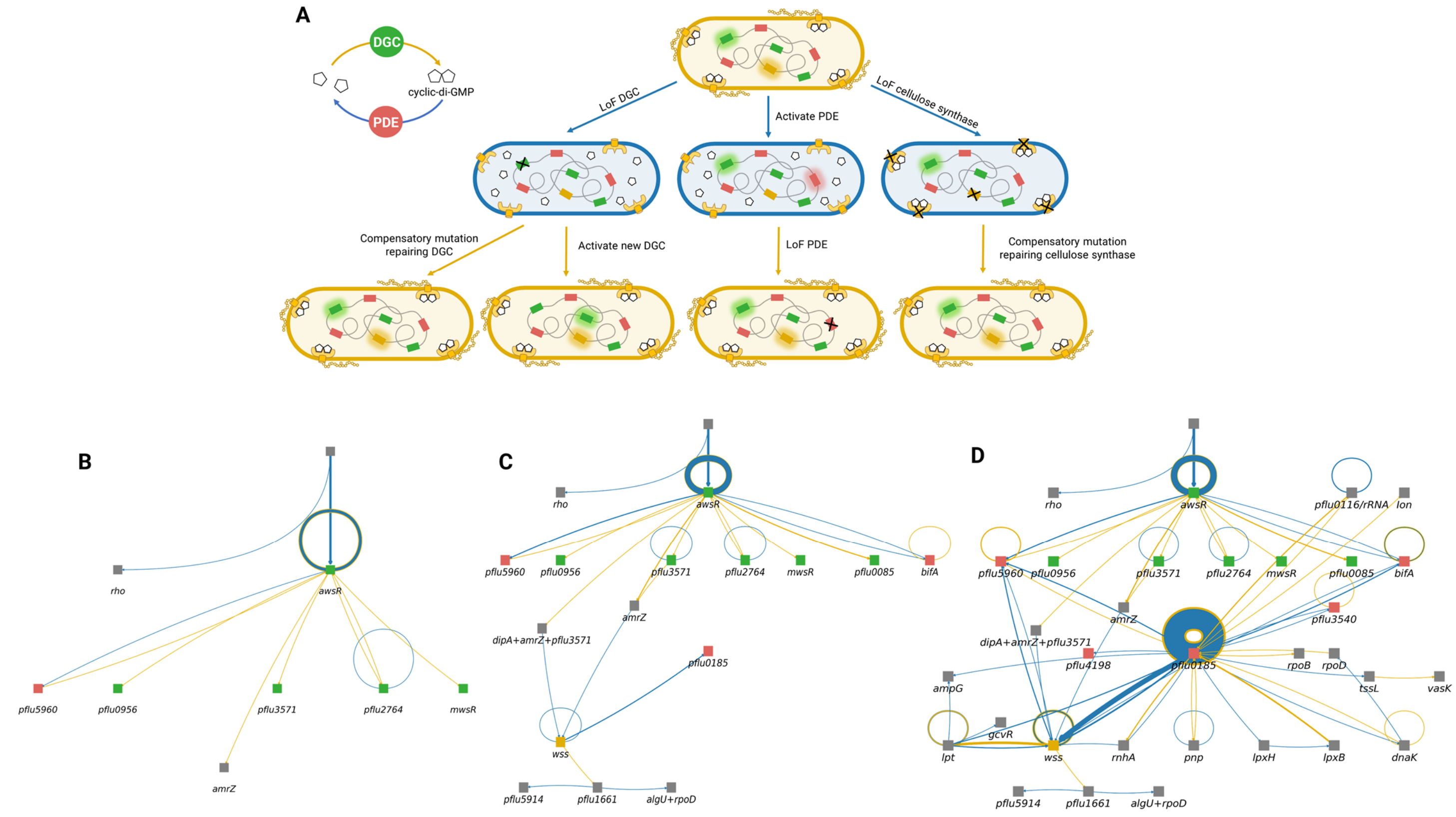


**Figure S3 Diversity of mutational paths generating variation in evolutionary potential.** (**A**). Genetic architecture underpinning successive mutational transitions between CEL$^{-}$ and CEL$^{+}$ states. Cyclic-di-GMP is the critical signalling molecule and allosteric activator of cellulose synthesis. The molecule is synthesised from two molecules of guanosine-5'-triphosphate (diamond symbols) by diguanylate cyclases (DGCs) and degraded by phosphodiesterases (PDEs), which function as positive and negative regulators, respectively. Red boxes in cells denote genes encoding PDEs, green boxes DGCs, and yellow box the structural determinant of cellulose biosynthesis encoded by the 10 gene *wss* operon. "Glow" indicates gene (or gene product) activation. The CEL$^{+}$ state is realised by mutations that elevate cyclic-di-GMP levels. Once in the CEL$^{+}$ state, there exist three primary routes to CEL$^{-}$: mutational inactivation of DGCs, mutational activation of PDEs, or LoF mutations in cellulose synthesis. The mutational trajectory a lineage takes affected its evolutionary potential. For example, lineages that transition to CEL$^{-}$ through destruction of the cellulose biosynthetic machinery were ~10 times more likely to go extinct at the next transition compared to any other target ($P < 0.0001$, Fisher's exact test, two-tailed). This stems from the fact that this machinery is essential for the $CEL^+$ and so must be repaired in the next transition to avoid extinction. In contrast, numerous regulatory genes controlling cyclic-di-GMP levels provided opportunity for mutation to successively activate and inactivate cellulose biosynthesis via redundant targets (the SBW25 genome encodes 41 genes with roles in synthesis and degradation of cyclic-di-GMP). Additional flexibility was evident in individual regulatory genes. For example, LoF caused by substitutions were often compensated by changes at residues elsewhere in the same regulatory gene (or through mutations in the 5'-untranslated region that likely increased gene expression). In contrast, not a single such instance was observed for the structural locus (*wss*), where repair came about almost exclusively through direct reversions (Supplementary Table S1). This indicates a much tighter structure-function relationship for the *wss* locus, contributing to this target being a frequent dead-end. The entire set of mutations and phenotypic effects can be explored at: micropop.evolbio.mpg.de/data/evolve. (**B-D**) Snap shots of mutational trajectories at transition T6 (**B**), T13 (**C**) and T41 (**D**) for all lineages from meta-population B starting from the ancestral state (the $CEL^+$ genotype used to found all eight lineages). The full sequence (up to T41) for all lineages is available as a video (Supplementary Movie S1), allowing mutations to be viewed transition-by-transition. Blue lines indicate transitions to $CEL^-$ and yellow lines transitions to $CEL^+$ as in Figure S3A. Boxes indicate genetic targets underpinning transitions. Line thickness indicates the frequency with which the mutational route was used. Self-loops, for example around *awsR*, indicate repeated transitions via successive activating / inactivating mutations. Extinction events are evident where a line enters but does not exit a node. (**B**) shows mutations in a range of genes, most of which are positive regulators of cellulose biosynthesis (DGCs, green squares), and one of which, the DGC-encoding *awsR*, was targeted by mutation multiple times, effecting repeated transitions between $CEL^-$ and $CEL^+$. By transition T13 (**C**), an increasing number of transitions were triggered by further cycles of mutation targeting *awsR*, however, utility of *awsR* ceased upon occurrence of a 54 base pair deletion (at T9 M6), which fixed within the meta-population at transition 12, following an almost fatal extinction event. Also notable at transition T13 are mutations that target negative regulators of cellulose biosynthesis (PDEs, red squares). Striking, however, is the fact that *pflu0185* became the target of multiple mutations effecting repeated switching between $CEL^-$ and $CEL^+$ states (thick self-loop circling *pflu0185*).

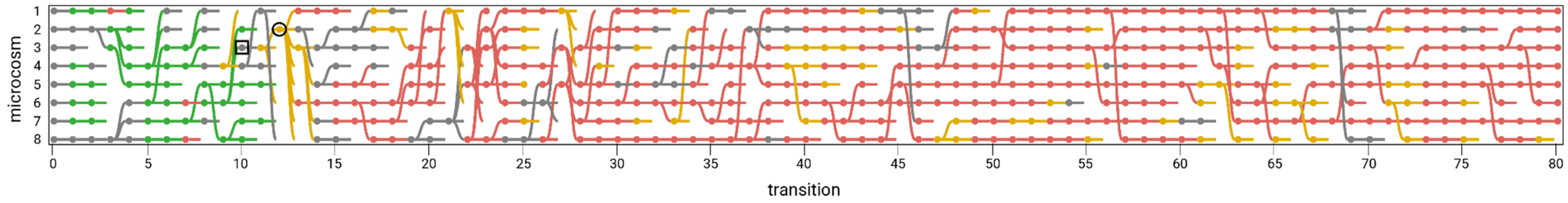


**Figure S4. Sudden and persistent change in the identity of regulatory targets due to mutation causing distributed production of cyclic-di-GMP.** The genealogy of meta-population B as shown in Figure 2B and 3B, but here nodes are coloured according to function of the transition-effecting mutation: green = DGC; red =PDE; yellow = cellulose synthesis; grey = other. Evident is the abrupt change in use of DGCs. At T10 M3 (black square), in the lineage that later went to fixation at T12 M2 (black circle), three atypical mutations co-occurred. While generation of the CEL$^{+}$ state had before this involved activation of a single DGC-encoding gene, these mutations targeted genes encoding a PDE (*dipA*), a DGC (*pflu3571*) and the global transcriptional regulator *amrZ*. The latter is conserved across *Pseudomonas* where it controls transcription of multiple DGCs [42]. We hypothesized that this unusual triple-mutant path had caused the source of cyclic di-GMP to become distributed across multiple DGC-encoding loci, such that LoF mutations at any single DGC, were no longer able to reduce cyclic-di-GMP levels sufficient to generate the CEL$^{-}$ state. In support of this hypothesis, transposon mutagenesis was employed in an attempt to identify genes capable of suppressing the phenotype of CEL$^{+}$ at T10 M3 (black square), but no DGC-encoding loci were identified. Assuming a Poisson distribution of transposon insertions across the SBW25 genome (6,700 genes), the probability of a gene remaining undetected given our screen of 54,000 mutants is ~0.0003. From transition 14 onward successful lineages no longer generated CEL$^{-}$ via changes in DGCs, but did so instead by mutations that affected the activity of PDEs. The shift from DGCs as targets, to PDEs, marked a significant contraction in the range of mutational targets and was instrumental in turning the focus of selection to negative regulators and increasingly to *pflu0185*.

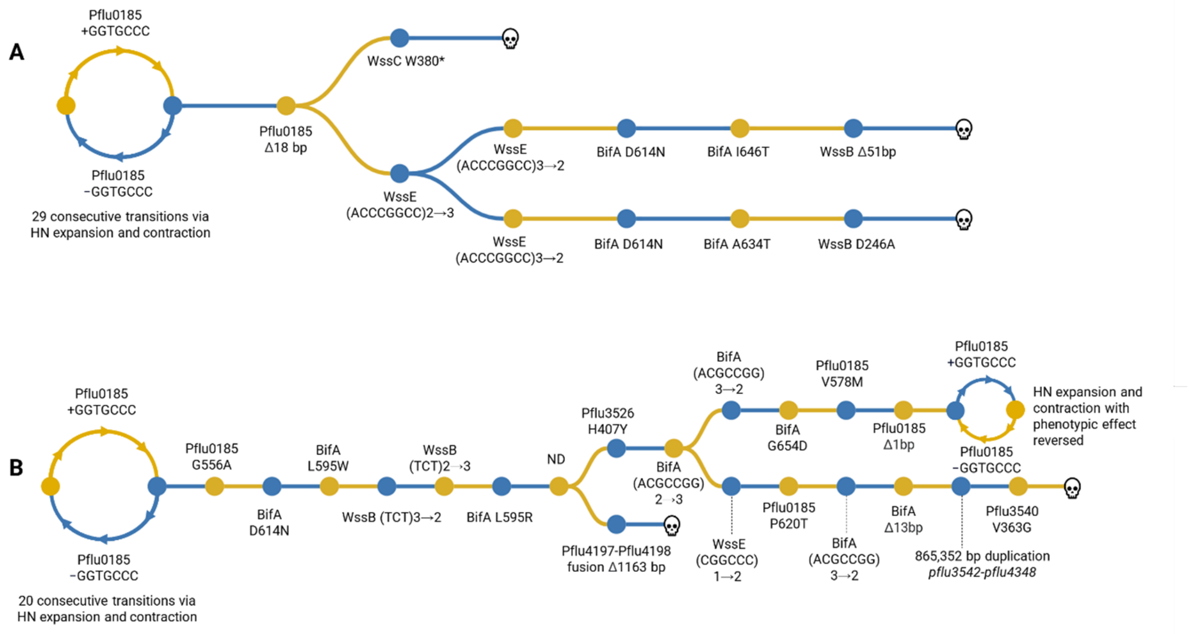


**Figure S5. Fates of lineages following alternative LoF mutations within *pflu0185*.** (**A**) Example of how an irreparable LoF mutation in *pflu0185* leads to lineage extinction. Following 29 successive transitions between CEL$^+$ and CEL$^-$ via changes in the number of HN repeats, this lineage acquires an 18 bp deletion in *pflu0185* as an alternative means of LoF. Despite loss of *pflu0185* this lineage leaves two offspring, both of which achieve (independently) CEL− by LoF mutations in the cellulose synthetic locus *wss*. Unsurprisingly, one faced extinction at the next transition, but the other, which became CEL$^-$ by duplication of an 8 bp sequence in *wssE*, left two offspring, with both achieving CEL$^+$ by exact reversion of the mutation at the next transition. The two extant lineages then transitioned to CEL$^-$ and then CEL$^+$ by changes to the PDE BifA, only to then acquire LoF mutations in *wssB*, followed by extinction. (**B**) Alternative LoF (G556A) mutation within *pflu0185* leads to trajectory through alternative PDEs (BifA, Pflu4198 and Pflu3540). The alternative *pflu0185* LoF mutation is in this case evidently reparable, with lineages returning to *pflu0185*. In one sub-lineage, HN expansion and contraction even restarts (upper branch), although this time with a reversed phenotypic effect. This is a consequence of a 1 bp LoF mutation occurring shortly downstream (17 bp) of the HN-sequence. Addition of 7 bp in this context restores the *pflu0185* reading frame. ND=not detected. The entire set of mutations and phenotypic effects can be explored at: micropop.evolbio.mpg.de/data/evolve

| ***wss* LoF mutation** | **Repair mutation** |
|---|---|
| *wssB* K226N AAG→AAT | direct reversion N226K |
| *wssE* 9bp deletion | *pflu1056 lptF* P306S CCC→TCC |
| | *pflu1057 lptG* V331E GTG→GAG |
| | *pflu1056 lptF* P306S CCC→TCC |
| | *pflu0884 lptH* T74P ACC→CCC |
| *wssB* Q271* CAG→TAG | *271S TAG→TCG |
| *wssA* 1bp deletion | 2bp deletion re-framing |
| *wssB* Q627* CAG→TAG | *627W TAG→TGG |
| *wssE* G7→G8 +G | direct reversion -G |
| *wssB* W383* TGG→TGA | direct reversion *383W |
| | direct reversion *383W |
| *wssC* 1 bp deletion | 11 bp deletion re-framing |
| *wssE* G7>G6 -G | direct reversion +G |
| *wssB* -TCT 3→2 | direct reversion +TCT |
| *wssB* -TCT 3→2 | direct reversion +TCT |
| *wssC* +TGCTGGTCAACG 1→2 | direct reversion - TGCTGGTCAACG |
| *wssE* +ACCCGGCC 2→3 | direct reversion - ACCCGGCC |
| | direct reversion - ACCCGGCC |

**Table S1.**

Loss-of-function mutations in genes determining the biosynthesis of cellulose and their means of repair.

| Locus | Gene | Events | Function |
|---|---|---|---|
| *pflu0491* | | 1 | outer membrane TolC efflux protein |
| *pflu0637* | *bfiS* | 1 | sensor kinase |
| *pflu1119* | | 1 | hypothetical protein |
| *pflu1372** | | 1 | hypothetical protein; polysaccharide deacetylase family |
| *pflu1479* | | 1 | metallo-β-lactamase |
| *pflu1593** | | 1 | hypothetical protein |
| *pflu1654* | | 1 | hypothetical protein; Glycosyl transferase family 2 domain |
| *pflu1657* | *fnl1* | 1 | polysaccharide biosynthesis protein |
| *pflu1687* | | 4 | methyl-accepting chemotaxis protein |
| *pflu1728** | *tpm* | 1 | thiopurine S-methyltransferase |
| *pflu1799** | *pyk* | 1 | pyruvate kinase |
| *pflu1815** | *gltA* | 1 | type II citrate synthase |
| *pflu1877/pflu1878* | | 1 | intergenic (-165/-748) |
| *pflu2181* | *rocS1* | 1 | two-component system sensor kinase |
| *pflu2227* | | 1 | putative homocysteine S-methyltransferase |
| *pflu2799/2800* | | 1 | intergenic (-141/-166) |
| *pflu3409* | | 1 | methyl-accepting chemotaxis protein |
| *pflu3835/pflu3836* | | 1 | intergenic (+43/+39) hypothetical proteins |
| *pflu3728** | | 1 | putative ribose ABC transporter ATP-binding protein |
| *pflu4184* | *cvpA* | 1 | colicin production membrane protein |
| *pflu4190* | *fimV* | 1 | motility regulator (flagella & type IV pili) |
| *pflu4395* | *atuD* | 1 | putative acyl-CoA dehydrogenase |
| *pflu4413* | *cheB* | 1 | chemotaxis-specific methylesterase |
| *pflu4418* | *fleN* | 2 | transcriptional regulator; FleQ antagonist |
| *pflu4551* | *aer* | 6 | aerotaxis receptor Aer |
| *pflu4593* | *xdhB* | 1 | xanthine dehydrogenase |
| *pflu4943* | | 1 | hypothetical membrane protein |
| *pflu5376* | | 1 | putative histone deacetylase family protein |
| *pflu5455** | | 1 | putative two-component system sensor kinase |
| *pflu5416** | *lipA* | 1 | lipoyl synthase |
| *pflu5729* | *dguC* | 1 | D-amino acid homeostasis |
| *pflu5739** | *mdcE* | 1 | malonate decarboxylase subunit gamma |
| *pflu5780* | *ftsY* | 1 | cell division protein |

**Table S2.**

Secondary mutations: All loci mutated exclusively alongside a known transition-mediating mutation.

| **Primary mutation (HN copy number)** | **Secondary mutation(s)** | **Function** |
| --- | --- | --- |
| 6→4 | *aer* Δ3bp | Aerotaxis receptor |
| 3→1 | *aer* S260P | |
| 2→1 | *aer* Δ9bp | |
| 2→1 | *aer* Δ21bp | |
| | *fimV* Q640* | Motility regulator (flagella and type IV pili) |
| 2→1 | *aer* E517K | Aerotaxis receptor |
| 2→1 | *aer* G270D | |
| 2→1 | *pflu1687* L27P | Methyl-accepting chemotaxis protein |
| 2→1 | *pflu1687* M154I | |
| 2→1 | *pflu1687* S6N | |
| 2→1 | *pflu1687* V215I | |
| 2→1 | *fleQ* R371C | Flagella transcription regulator |
| 2→1 | *fleN* L42Q | Flagella transcription regulator |
| 2→1 | *cheB* T92P | Chemotaxis-specific methylesterase |

**Table S3.**

Secondary adaptive mutations associated with changes in number of HN repeats

| | Locus | Mutation | Gene name | Function |
|---|---|---|---|---|
| **A** | *pflu0185* | N160S (AAC→AGC) | | PDE |
| | *pflu0185* | R175C (CGT→TGT) | | PDE |
| | *pflu0458* | F624L (TTT→CTT) | *dipA* | PDE |
| | *pflu4197/4198* | Δ932 bp fusion | | putative LysR family transcriptional regulator/PDE |
| | *pflu4439* | (TGG)2→4 coding (991-996/1782 nt) | *Flif* | flagellar M-ring protein |
| | *pflu4744* | L79P (CTG→CCG) | *AmrZ* | transcription factor cellulose synthase operon |
| | *pflu5210* | T366I (ACC→ATC) | *awsR* | DGC |
| | *pflu5211* | Δ33 bp coding (223-255/573 nt) | *awsX* | negative regulator AwsR |
| | *pflu5329* | Δ9 bp coding (3071-3079/3852 nt) | *mwsR* | PDE-DGC |
| | *pflu5329* | (GAGCTG)2→1 coding (3448-3453/3852 nt) | *mwsR* | PDE-DGC |
| | *pflu5960* | P358S (CCG→TCG) | | PDE |
| | *pflu5960* | Δ15 bp coding (819-833/1665 nt) | | PDE |
| **B** | *pflu0185* | upstream -53bp A→G | | PDE |
| | *pflu0185* | I158S (ATC→AGC) | | PDE |
| | *pflu0414* | V550A (GTG→GCG) | *gltB* | glutamate synthase subunit alpha |
| | *pflu1224* | (GCG)4→3 coding (937-939/1011 nt) | *wspF* | negative regulator of DGC WspR |
| | *pflu1225* | Δ234 bp coding (656-889/1002 nt) | *wspR* | DGC |
| | *pflu3027* | (GGC)2→1 coding (235-237/768 nt) | | putative short-chain dehydrogenase family protein CDS |
| | *pflu5210* | T27P (ACC→CCC) | *awsR* | DGC |
| | *pflu5329* | Δ9 bp coding (3071-3079/3852 nt) | *mwsR* | PDE-DGC |
| | *pflu5329* | (GAGCTG)2→1 coding (3448-3453/3852 nt) | *mwsR* | PDE-DGC |
| | *pflu5691* | R180L (CGC→CTC) | *dbpA* | ATP-dependent RNA helicase |
| **C** | *pflu1219* | Δ64 bp pflu1219 wspA 1362-1421 | *wspA* | Wsp pathway |
| | *pflu1224* | Δ3 bp residue 313 | *wspF* | negative regulator of DGC WspR |
| | *pflu1768* | G67G (GGC→GGA) | | hypothetical protein |
| | *pflu3027* | G88C (GGC→TGC) | | putative short-chain dehydrogenase family protein CDS |
| | *pflu5211* | Y57C (TAT→TGT) | *awsX* | negative regulator of DGC AwsR |
| **D** | *pflu1219* | F244S (TTC→TCC) | *wspA* | Wsp pathway |
| | *pflu1219* | A420V (GCC→GTC) | *wspA* | Wsp pathway |
| | *pflu1223* | R302C (CGC→TGC) | *wspE* | Wsp pathway |
| | *pflu1223* | A585V (GCC→GTC) | *wspE* | Wsp pathway |
| | *pflu4858* | F333L (TTC→CTC) | *bifA* | PDE |
| | *pflu4858* | M555I (ATG→ATA) | *bifA* | PDE |
| | *pflu5210* | T27P (ACC→CCC) | *awsR* | DGC |
| | *pflu5329* | Δ3 bp coding (2932-2934/3852 nt) | *mwsR* | PDE-DGC |
| | *pflu5329* | Δ9 bp coding (3071-3079/3852 nt) | *mwsR* | PDE-DGC |

**Table S4.**

Mutations in $CEL^+$ genotypes founding meta-populations A, B, C and D

**Movie S1.**

Sequential accumulation of mutations and genetic targets (up to T41) for all lineages of meta-population B viewed transition-by-transition. Nodes indicate genetic targets; green = DGC, red = PDE, yellow = cellulose biosynthesis machinery, grey = other. Line thickness indicates the frequency with which the mutational route was used. Blue lines indicate transitions to $CEL^{-}$ and yellow lines transitions to $CEL^{+}$ as in Figure 5A. Self-loops, for example around *awsR* and *pflu0185*, indicate repeated transitions via successive activating/inactivating mutations at these loci. Extinction events occur where a line enters but does not exit a node.
